# Comparing morphological and molecular methods for stream macroinvertebrate biodiversity assessment across seven watersheds in the southern United States

**DOI:** 10.64898/2026.09.24.754159

**Authors:** Daniel C. Allen, Zacchaeus G. Compson, Chelsea Smith, Michael T. Bogan, Lindsey E. Vande Streek, Kaley Cave, Arial Shogren, Carla L. Atkinson, Brian A. Gill, Luiza Gonçalves Lazzaro, Fagbohun Ibrahim, Meryl C. Mims, Travis Apgar, Albert Ruhi, Sean Emmons, Kierstyn T. Higgins, Stephen C. Cook, Samuel Silknetter, Kyle Leathers, Megan Malish, Michelle H. Busch, Thomas Neeson, Veronica Saenz, Rose Mohammadi, Yang Hong

## Abstract

Freshwater biodiversity is declining at alarming rates globally, yet our capacity to monitor it depends heavily on the methods used for biodiversity assessment. Stream benthic macroinvertebrates are widely used as bioindicators, but traditional morphological identification is labor-intensive and limited by taxonomic expertise, while DNA metabarcoding offers a promising but incompletely evaluated alternative. Here, we compared biodiversity patterns detected by four macroinvertebrate sampling and identification methods (morphological identification of benthic samples, morphological identification of combined benthic and stream-edge samples, eDNA metabarcoding of bulk benthic samples, and eDNA metabarcoding of streamwater) across 62 sites in seven watersheds spanning the southern continental United States. We compared methods across three facets of biodiversity (gamma diversity, alpha diversity, and taxonomic composition) and three taxonomic levels (family, genus, and species). We found that which method detected more taxa shifted with taxonomic resolution: morphological methods detected more families and genera, while metabarcoding methods detected more species. Bulk benthic metabarcoding detected more families and genera than water eDNA metabarcoding, consistent with the “watered-down biodiversity” effect of DNA dilution in water samples, but the two metabarcoding methods did not differ significantly in species richness. All four methods detected largely distinct suites of taxa, with metabarcoding methods yielding more unique taxa than morphological methods. Notably, taxa uniquely detected by water eDNA were dominated by soft-bodied organisms (oligochaetes, leeches, earthworms) and water-column-associated taxa, while benthic eDNA uniquely detected hard-bodied EPT insects. We attribute this pattern to fundamental differences in eDNA shedding, transport, and deposition dynamics across organism types. These results demonstrate that no single method captures a complete picture of stream macroinvertebrate biodiversity, and that method choice should be guided by the taxonomic resolution and community components most relevant to the goals of the bioassessment program.

## Introduction

Freshwater ecosystems harbor a disproportionate share of the planet’s biodiversity relative to their spatial extent, and are experiencing the steepest rates of biodiversity loss of any biome on Earth (Dudgeon and Strayer 2025). Global monitoring efforts indicate that freshwater vertebrate populations have declined by roughly 85% since 1970, more than 50% greater than the rate of decline recorded in terrestrial or marine systems, and nearly one-quarter of assessed freshwater species are now threatened with extinction (WWF 2024). These declines are driven by habitat fragmentation and flow modification by dams, pollution, invasive species, and climate change, often acting in combination (Dudgeon et al. 2006, Reid et al. 2019). Reversing or even slowing these trends will require the ability to detect and track changes in freshwater biodiversity across broad spatial and temporal scales, yet our capacity to do so remains fundamentally constrained by the methods available for biodiversity assessment (Simaika et al. 2024). Choosing among these methods may influence which taxa are detected, how diversity patterns are characterized, and ultimately, what conservation and management conclusions are drawn from monitoring data.

Among freshwater taxa, benthic macroinvertebrates are uniquely well-suited for biodiversity assessment and have long served as the backbone of stream biomonitoring programs (Rosenberg and Resh 1993, Barbour et al. 1999). Their ubiquity across stream types, relatively sedentary life histories, and well-characterized sensitivities to environmental stressors make them sensitive and interpretable indicators of water quality and biotic integrity (Wallace and Webster 1996, Bonada et al. 2006). Decades of research have linked macroinvertebrate community composition to specific stressors, including nutrient enrichment, sedimentation, and flow alteration, providing the foundation for widely used biotic indices and rapid bioassessment protocols implemented by state and federal agencies (Karr 1981, Hilsenhoff 1987, Barbour et al. 1999). However, traditional morphological identification of macroinvertebrates is labor-intensive, requires substantial taxonomic expertise, and often resolves only coarse taxonomic levels for certain taxa, limiting the spatial and temporal scope over which bioassessment can realistically be conducted.

Environmental DNA (eDNA) metabarcoding has emerged as a promising alternative or complement to traditional morphological identification, offering the potential to standardize and accelerate macroinvertebrate biodiversity assessment across large spatial scales (Baird and Hajibabaei 2012). By sequencing genetic material present in benthic samples or suspended in streamwater, metabarcoding can simultaneously detect diverse taxa without the time and expertise demands of morphological sorting and identification, and without reliance on the presence of identifiable life stages (Carew et al. 2013). These features make eDNA-based approaches particularly attractive for large-scale and long-term monitoring programs, where consistent, efficient, and reproducible sampling across many sites is essential (Leese et al. 2018). Indeed, metabarcoding has been increasingly adopted across freshwater systems for biodiversity surveys (Deiner et al. 2017), invasive species detection (Comtet et al. 2015), and rare species monitoring (Smart et al. 2015), with proponents arguing that it could eventually standardize bioassessment protocols across regions and agencies that currently rely on disparate taxonomic and methodological traditions (Leese et al. 2018, Bohmann et al. 2021).

Despite this promise, we still lack large-scale, multi-site studies that rigorously compare the biodiversity patterns recovered by traditional morphological identification versus eDNA metabarcoding for stream macroinvertebrates (but see (Emmons et al. 2023). Most existing comparisons are limited to a small number of sites or a single watershed, making it difficult to evaluate whether observed differences between methods are generalizable or idiosyncratic to local conditions (Elbrecht et al. 2017, Emmons et al. 2023). Moreover, within eDNA-based approaches, metabarcoding can be performed on bulk benthic samples (capturing organismal DNA from sediment and detritus) or on water samples (capturing eDNA shed or suspended in the water column), and these two sample types differ in the taxa they detect due to differences in DNA source, transport, and degradation (Deiner et al. 2016, Hajibabaei et al. 2019a). Few studies have directly compared benthic-sample and water-sample metabarcoding against traditional morphological identification within the same set of sites across a large spatial scale, leaving open key questions about whether these approaches are interchangeable, complementary, or fundamentally divergent in the biodiversity signal they capture.

Here, we leverage a large-scale field effort across seven watersheds spanning the southern continental United States to directly compare biodiversity patterns recovered by four macroinvertebrate sampling and identification approaches: traditional morphological identification of benthic samples, morphological identification of combined benthic and stream-edge samples (i.e., riparian microhabitats such as roots, partially flooded vegetation, and pooled backwaters; not typically included in benthic biomonitoring), DNA metabarcoding of benthic samples, and eDNA metabarcoding of streamwater samples. Using paired samples collected at the same sites and times, we compare these four methods across multiple facets of biodiversity, including gamma diversity, alpha diversity, and taxonomic composition, and across multiple levels of taxonomic resolution (family, genus, and species). We hypothesize that metabarcoding and morphological methods will converge in estimates of broad taxonomic diversity (e.g., order, family) but diverge at finer taxonomic resolution (genus, species), reflecting differences in reference database coverage and taxonomic resolving power between methods. We further hypothesize that benthic and water-based metabarcoding will detect distinct subsets of taxa due to differences in DNA source and transport, and that no single method will fully capture the macroinvertebrate biodiversity recovered when all methods are considered together. By identifying where these methods converge and diverge, this study provides critical guidance for researchers and monitoring programs seeking to incorporate eDNA-based approaches into large-scale freshwater biodiversity assessment.

## Methods

### Field Sampling

We established study sites at streams across seven watersheds spanning the southern half of the continental US (Figure 1). Ten candidate stream sites were selected in each watershed, and were sampled in 2022, in the season when streams were most likely to be wetted. These watersheds were selected as part of the broader StreamCLIMES project that investigated how drying impacts stream benthic macroinvertebrate communities (Gill et al. 2024). The 10 stream sites were selected to encompass a range of stream habitats and various drying frequencies. Because some sites were completely dry in some years but not others, not all streams were wetted and thus sampleable in 2022. We present data from 62 sites sampled across these seven basins (Table S1).

**Figure 1.**
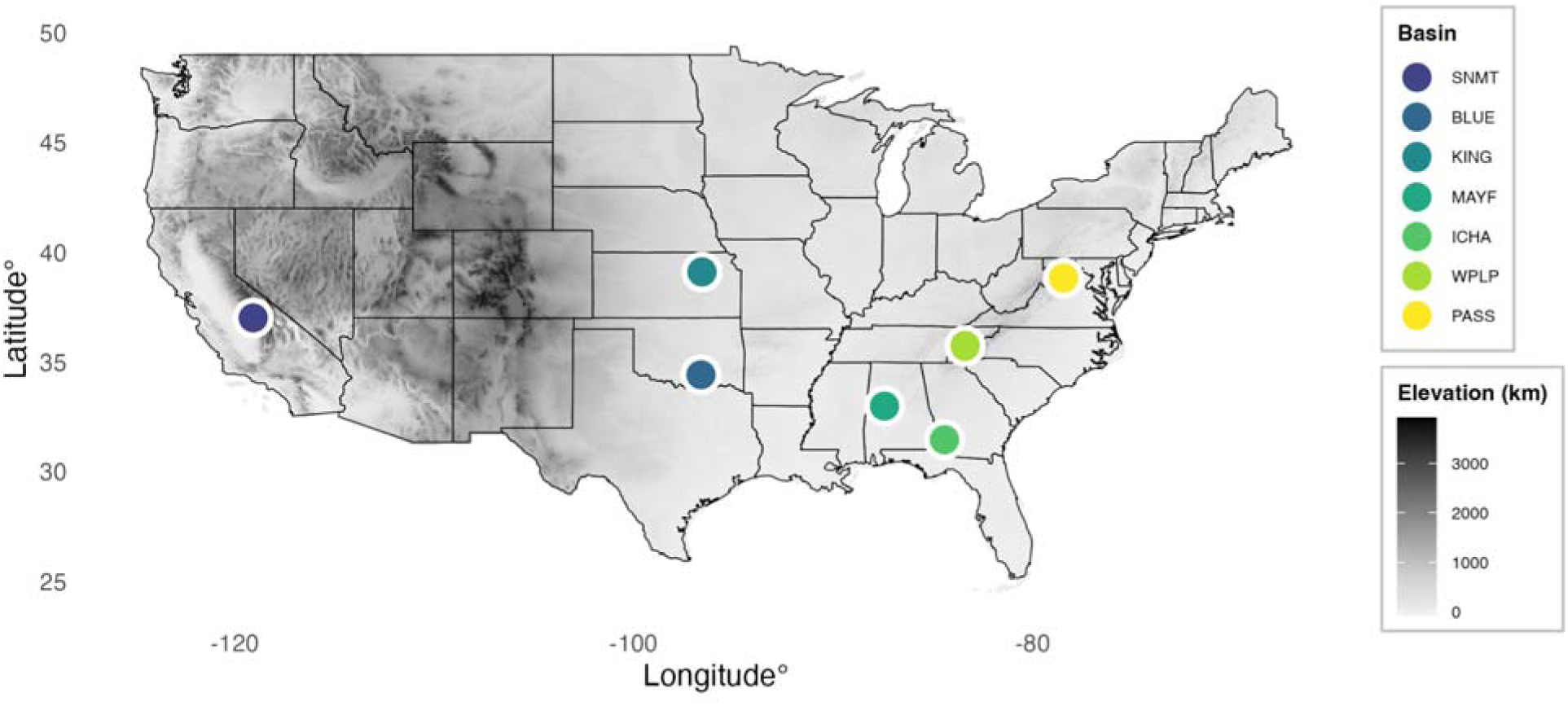
US elevation map (WGS84 projection) showing the locations of the stream basins used in this study. Basin name abbreviations: SNMT, Sierra Nevada Mountain Tributaries; BLUE, Blue River; KING, Kings Creek; MAYF, Mayfield Creek; ICHA, Ichawanochoway Creek; WPLP, West Prong Little Pigeon River; PASS, Passage Creek.

We collected benthic macroinvertebrate samples at each study site following (Gill et al. 2024). Briefly, we established a 150 m study reach at each site with transects every 15 meters along the reach, resulting in eleven transects total. When possible, we used a Surber sampler with a 500 μm mesh net to sample benthic macroinvertebrates at the left bank, center, or right-bank position along each transect, alternating positions along each transect. In cases with deeper or non-flowing water, a D-frame net was used to sample the same benthic area. The eleven samples (0.0929 m2 each) were combined into a composite that comprised a total sample area of 1.02 m2. In some cases, the total area sampled was lower if dry reaches prevented sampling at some transects but not others, or if low water levels required mini-surber samplers (0.0232 m² each); see (Gill et al. 2024) for more details. Additionally, we collected a composite sample of five sweeps (∼□0.33□m2 each) along each reach’s wetted edge in riparian-adjacent microhabitats as an edge sample with a 500 μm mesh D-frame net. Edge samples were processed and identified by the same personnel at the University of Arizona.

For the purposes of this study, we collected duplicate samples simultaneously, such that the first could be used for traditional morphological identification and the second could be used for DNA metabarcoding of the bulk sample. Sampling equipment was treated with 50% bleach solution to prevent DNA contamination across sites (Bustin et al. 2009, Goldberg et al. 2016). We sent the first set of benthic samples to the University of Arizona to process for traditional morphological identification, and the second set to Penn State University for processing for DNA metabarcoding (lab methods described below).

Prior to any sampling or disturbance in the stream made by the sampling crew, we also collected triplicate 1 L water samples at the bottom of the 150 m study reach for collection of environmental DNA suspended in streamwater. We collected water samples as close to the thalweg (the deepest part of the channel) as possible. We kept water samples on ice, and immediately froze them after returning from the field site later that day. We shipped frozen water samples to the University of North Texas where they were stored at −80 C until they were processed.

### Sample Processing and DNA Extraction

Benthic and edge samples for morphological identification were processed in the laboratory following (Gill et al. 2024). A minimum of 500 individuals were identified per sample, splitting the sample as necessary. We then identified specimens to at least genus-level for insects and to order-or family-for non-insects using available keys, and to species when possible (Cook 1974, Merritt et al. 2008, Andersen et al. 2013, Thorp and Rogers 2015). We used benthic and edge sample data to generate two different data sets: a “benthic only” sample that did not include edge sample data and a “benthic + edge” dataset that included sample data from both methods. Because the edge samples were qualitative, while the benthic samples were quantitative, our “benthic + edge” dataset used the approach recommended by (Gill et al. 2024) where any taxon present in the edge but not the benthic sample was given an abundance of 1.

Benthic samples for DNA metabarcoding were stored in 95% EtOH at −20 C prior to being processed. We removed pieces of large debris (e.g., sticks and leaves) prior to homogenization. Sample remainders were homogenized using a commercial blender for 2-3 minutes on ice, before transferring 50 ml of the homogenate to a conical tube and centrifuging for 20 minutes at 7830 RPM in a vacufuge. Blenders were treated with 50% bleach solution to avoid cross-contamination of samples (Bustin et al. 2009, Goldberg et al. 2016). After the initial centrifugation, the supernatant was removed, and the remainder was vacufuged until the samples were fully dry. DNA was then extracted from 175 g of the dried homogenate using QIAGEN PowerSoil Pro DNA extraction kits. For water samples processed for eDNA metabarcoding, samples were thawed and filtered using 0.45 μm MCE (Nitrocellulose Mixed Ester) filters, and DNA was extracted from filters using Qiagen DNeasy Blood and Tissue extraction kits with an extended 48-hour incubation step to maximize DNA yield (see (Carim et al. 2016).

### DNA Metabarcoding

Extracted DNA from both benthic and water samples were sent to Jonah Ventures (Boulder, CO) for metabarcoding. We used three CO1 primers for PCR amplification to capture a wide range of invertebrate taxa: F230, BR5, and ml-jg (Hajibabaei et al. 2012, 2019b, Leray et al. 2013). PCRs were run with initial denaturation at 95 C for 5 minutes, followed by 40 cycles of 40 seconds at 95 C, 1 minute at 46 C, 30 seconds at 72 C and a final elongation at 72 C for 10 minutes. Following pooling and normalization, sample libraries were sequenced on an Illumina NovaSeq 6000 (San Diego, CA) at the Texas A&M AgriLife Genomics and Bioinformatics Sequencing Core facility using the SP Reagent Kit v1.5. Amplicons were sequenced in two sequencing runs, the benthic sample and water samples were sequenced separately.

### Bioinformatics

Raw sequence data were demultiplexed using pheniqs v2.1.0 (Galanti et al. 2021), enforcing strict matching of sample barcode indices (i.e, no errors). Cutadapt v3.4 (Martin 2011) was used to remove gene primers from the forward and reverse reads, discarding any read pairs where one or both primers were not found at the expected location (5’) with an error rate < 0.15. Read pairs were then merged using vsearch v2.15.2 (Rognes et al. 2016), discarding resulting sequences with a length of < 219 bp, > 239 bp, or with a maximum expected error rate > 0.5 bp (Edgar and Flyvbjerg 2015). For each sample, reads were then clustered using the unoise3 denoising algorithm (Edgar 2016) as implemented in vsearch, using an alpha value of 5 and discarding unique raw sequences observed less than 8 times. Counts of the resulting exact sequence variants (ESVs) were then compiled and putative chimeras were removed using the uchime3 algorithm in vsearch. For each final ESV, a consensus taxonomy was assigned using a custom best-hits algorithm and a reference database consisting of publicly available sequences (GenBank (Benson et al. 2005)) as well as Jonah Ventures’ in-house voucher sequence records. Reference database searching used an exhaustive, semi-global, pairwise alignment with vsearch, and match quality was quantified using a custom, query-centric approach, where the % match ignored terminal gaps in the target sequence, but not the query sequence. The consensus taxonomy was then generated using either all 100% matching reference sequences or all reference sequences within 1% of the top match, accepting the reference taxonomy for any taxonomic level with > 90% agreement across the top hits.

### Data Cleaning

We cleaned the taxonomy data from Jonah Ventures by sequentially filtering reads based on a conservative percent confidence match: order-level: reads ≤ 90% were removed; family-level reads ≤ 95% were removed; genus-level reads ≤ 98% were removed; and species-level reads ≤ 99% were removed (Lanzén et al. 2012, Pappalardo et al. 2021). We then removed all terrestrial taxa and non-invertebrate taxa (e.g., terrestrial ants, algal taxa, fungal taxa) by inspecting the remaining taxa. Next, we consolidated the primer taxa-site list to form a single dataset with the presence/absence of taxa across sites.To increase compatibility with morphological data, undescribed species (e.g., Simulium sp. BOLD:AAA9710) were combined with genus-level identifications, followed by subsequent checking of each sample to remove any higher-level taxa that were present and identified to a lower level (e.g., Chironomidae family removed if Chironomus genus present in sample).

### Data Analysis

We first examined gamma diversity to assess broad patterns of diversity across the four sampling methods, using the subset of 42 sites where we had data from all four methods—Morphological-Benthic, Morphological-Benthic+Edge, Metabarcoding-Benthic, and Metabarcoding-Water. Gamma diversity was calculated as the total number of taxa identified at each taxonomic level (Phylum, Class, Order, Family, Genus, and Species) for each dataset. To calculate gamma diversity, datasets from the four methods were processed using the dplyr and tidyr R packages (Wickham et al. 2019). This and all subsequent analyses were conducted using R vs. 4.5.1 (R Core Team 2025).

We also used taxa accumulation curves to assess richness patterns across genomic and morphological methods, again restricting this comparison to the sites where each method was used. Abundance data from morphological methods and read data from DNA metabarcoding methods were processed into presence-absence matrices. Accumulation curves were computed for each method using the specaccum function in the vegan package (Oksanen et al. 2025), and results were visualized using the tidyverse suite of packages (Wickham et al. 2019).

We compared alpha diversity detected by different bioassessment methods in two ways. First, we used the log-response-ratio approach described by (Hedges et al. 1999). We calculated the log response ratio at each site as ln(A/B), where A represents the total alpha diversity detected by one method (e.g. benthic sample DNA metabarcoding) and B represents the alpha diversity detected by another (e.g. benthic sample morphological identification). For alpha diversity, we made the following methodological comparisons: 1) benthic sample DNA metabarcoding vs. benthic sample morphological identification, 2) water eDNA metabarcoding vs. benthic + edge sample morphological identification, and 3) benthic sample DNA metabarcoding vs. water eDNA metabarcoding; these comparisons were represented at the family, genus, and species taxonomic levels. We assessed if log-ratios significantly deviated from zero using the LmerTest package in R (Kuznetsova et al. 2017) to assess a linear mixed model with watershed as a random effect to account for potential differences among watersheds generated using the lme4 package (Bates et al. 2015). Second, we performed linear regressions to compare taxonomic richness obtained by each method across taxonomic levels. For each pairwise comparison, we used what we hypothesized to be the most widely used method as the predictor variable and the least widely used method as the response variable (in decreasing order: morphological identification, DNA metabarcoding of homogenized benthic samples, and eDNA metabarcoding of streamwater).

Finally, we compared the taxonomic compositions of the macroinvertebrate community datasets produced by the different methods across taxonomic levels. Because of the uncertainty regarding how well abundance estimates obtained by visual enumeration in morphological identification relate to read abundances obtained by DNA (Lamb et al. 2019, Luo et al. 2023), we used presence-absence data for non-metric multidimensional scaling (NMDS) ordinations and alluvial plots. We restricted these comparisons to the 42 sites where all four methods were used, and to reduce the influence of rare taxa, we removed those with a relative abundance of < 1% for a given site-method-taxonomic level combination (for abundance data, we used counts for morphological identifications and read abundances for metabarcoding identifications).

Because the species community matrix was too sparse (97.96% zeroes) for reliable ordinations and meaningful alluvial plots, we only present results for family and genus. For the NMDS, we used Jaccard distances and first attempted a 2-axis solution (with try = 200 and trymax = 1,000). If the stress value of the 2-axis solution > 0.2, or if it did not converge, we then attempted a 3-axis solution, which was required at the family level. We present visualization of the first two axes here and visualizations of all three axes in supplemental information (Figure S1). For genus level analyses, NMDS ordinations revealed a severe outlier (4315% greater than the next highest value), so this site (MAYF-10) was removed from the dataset. To visualize differences in taxonomic compositions across family, genus, and species levels, we used alluvial plots (aka “Sankey diagrams”). We used the vegan and ggalluvial packages in R to perform the NMDS and to create alluvial plots, respectively. To assess differences in community composition between methods, we used the multi-response permutation procedure (MRPP).

## Results

### Gamma diversity

For gamma diversity, we found that DNA metabarcoding and morphological methods tended to be more similar at the phylum, class, order and family taxonomic levels for the total number of taxa identified (Figure 2). However, morphological methods appeared to identify more genera while DNA metabarcoding methods seemed to identify more species (Figure 2). Taxonomic accumulation curves mirrored these trends, showing greater diversity accumulation on a per sample basis for morphological approaches at family and genus levels. Interestingly, the rate of accumulation at the family level for morphological samples appeared to be approaching their maxima, while benthic-metabarcoding showed potential for increased accumulation with more sampling. At the species level greater diversity within DNA metabarcoding methods was seen, similar to previous trends (Figure 3).

**Figure 2.**
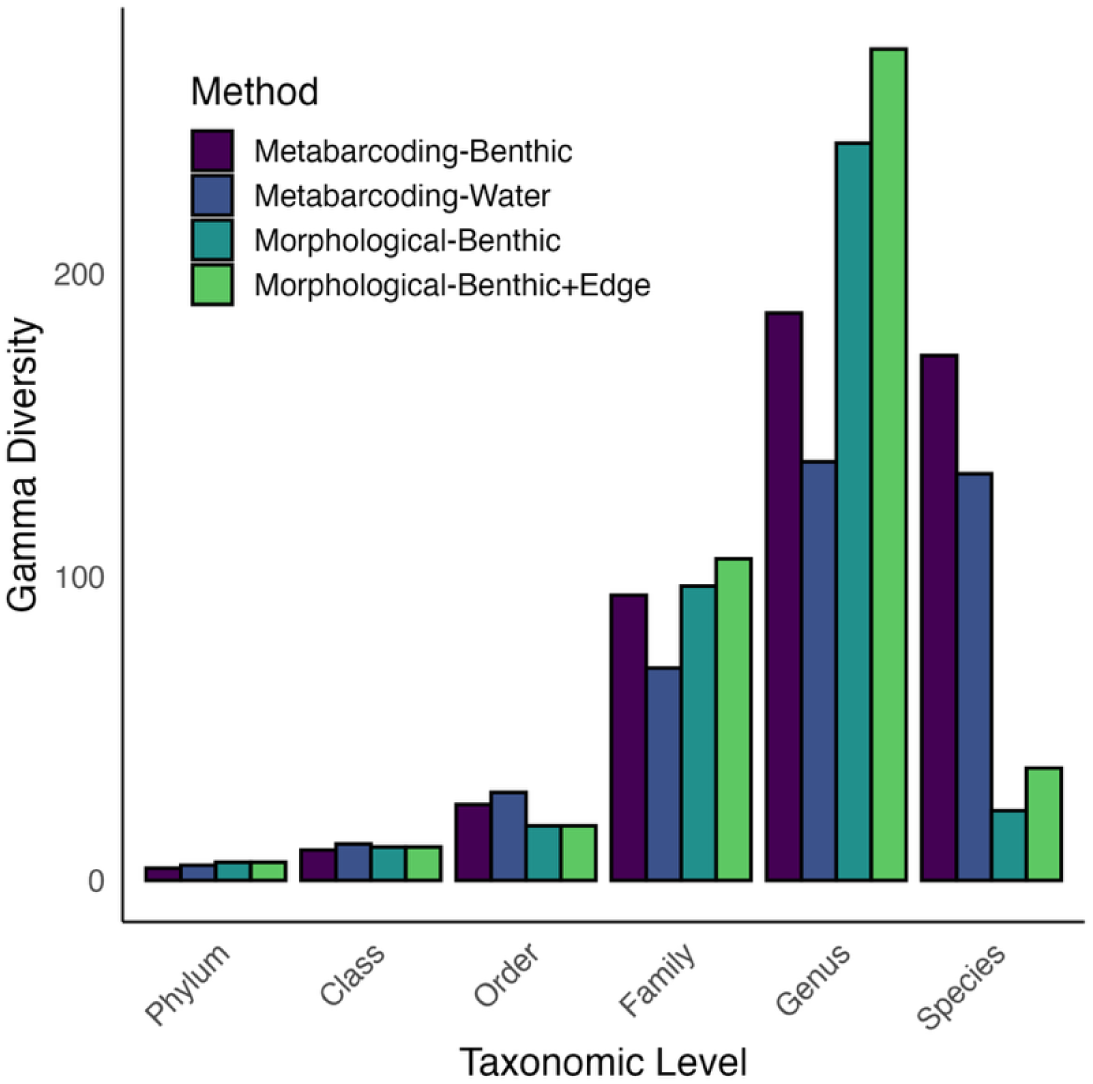
Gamma diversity bar plot of all basins indicating the total number of species for several different sampling methods, including those using DNA-based (“Metabarcoding”) and traditional (“Morphological”) approaches.

**Figure 3.**
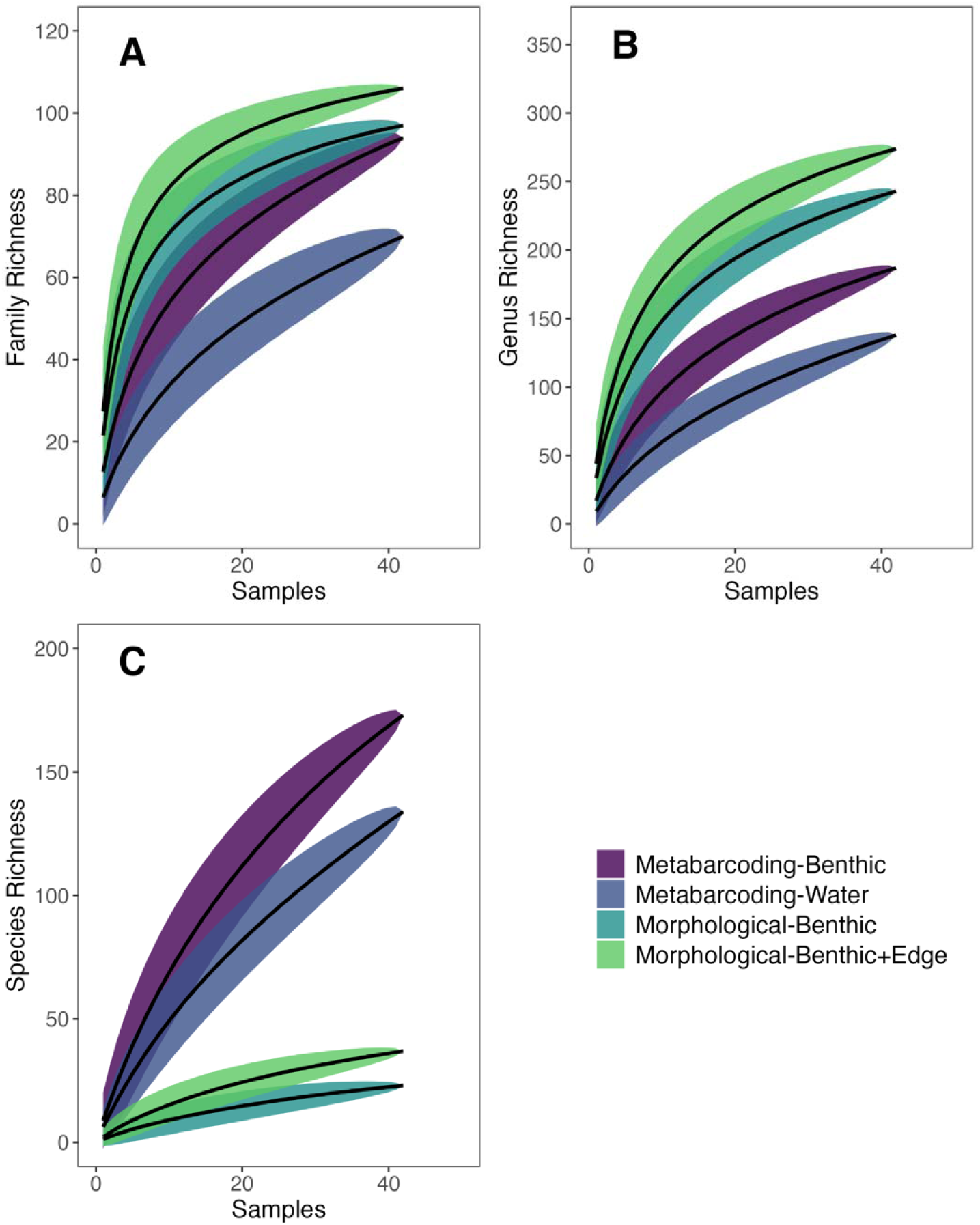
Accumulation curves displaying unique taxa richness with increasing sampling effort at Family (A), Genus (B) and Species (C) taxonomic levels for the four methods. Shaded areas show ±2 standard deviations around the mean estimates.

### Alpha Diversity

When comparing alpha diversity detected by the metabarcoding-benthic and the morphological-benthic methods, we found negative log-ratio values for family and genus, indicating that the morphological method detected more diversity than the metabarcoding method at these taxonomic levels. Positive log-ratios were recorded for species (Table 1, Figure 4), indicating that the morphological method detected more diversity than the metabarcoding method at these taxonomic levels. The linear mixed model regression analysis showed significant positive relationships of richness taxonomic levels between both methods (Table 2, Figure 5). When comparing water eDNA metabarcoding and morphological benthic plus edge methods, we observed negative mean log ratio values for family and genus, though a positive mean log-ratio value was observed at the species level (Table 1, Figure 4). The linear mixed model regression analysis also showed significant positive relationships of richness taxonomic levels between both methods (Table 2, Figure 5). Finally, when comparing water and benthic sample eDNA metabarcoding methods, we found significant negative log ratios for family and genus, but the log ratio for species was not significantly different from zero (Table 1, Figure 4). Moreover, linear mixed model regressions showed a significant relationship between family richness values of both methods, but not for genus or species richness (Table 2, Figure 5).

**Figure 4.**
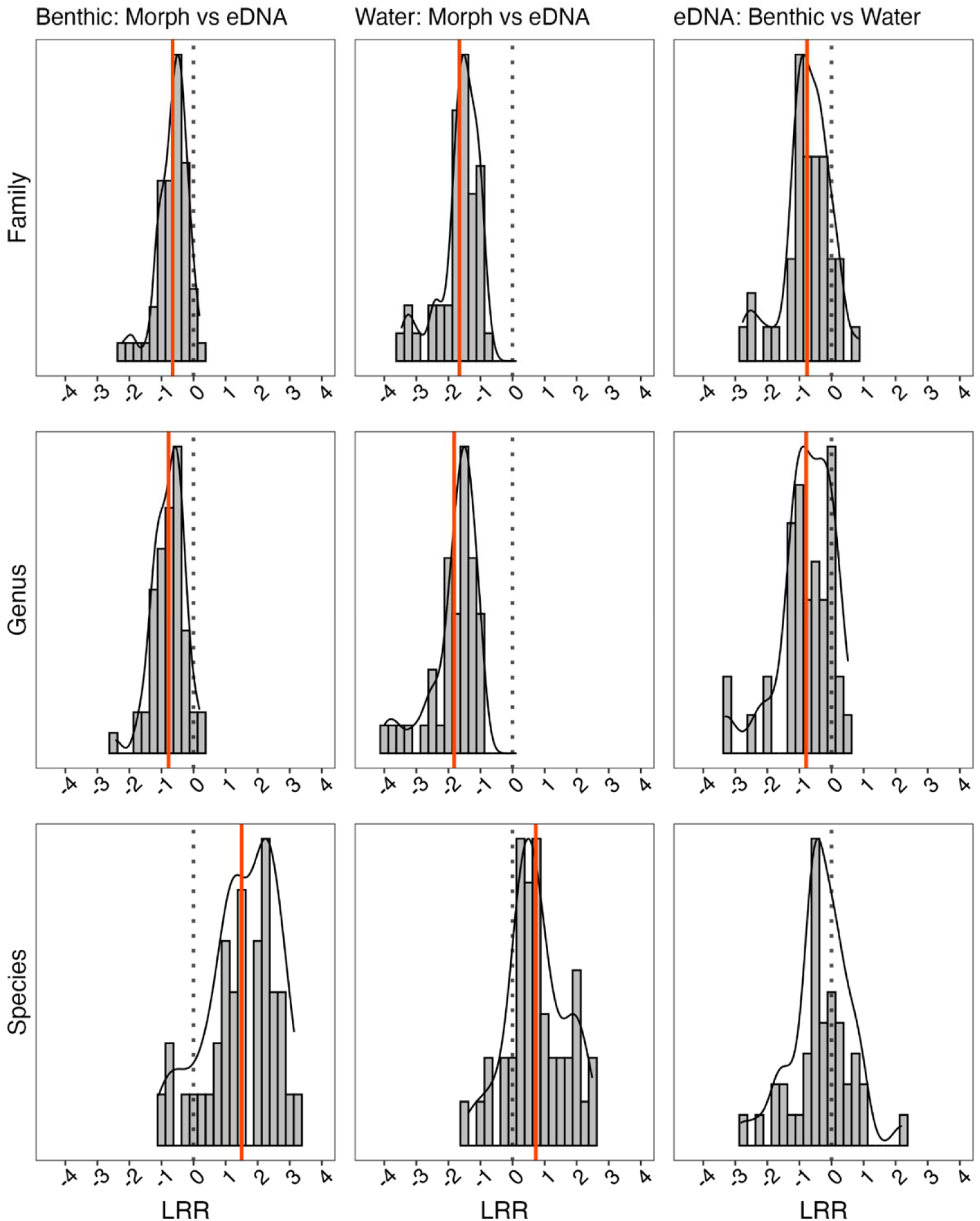
Histograms showing the log response ratios (LRR) of alpha richness detected by different identification methods. In graph labels, “A vs B” indicates LRR of ln(B/A), such that negative values indicate method B identifies fewer taxa. Panels correspond to a pairwise comparison of two methods at a given taxonomic level. Dotted lines indicate LRR values of zero, and the red lines represent mean LRR values that are statistically significant from zero. Left column: DNA metabarcoding benthic compared to morphological benthic samples. Middle column: morphological benthic + edge samples compared to eDNA metabarcoding of streamwater. Right column: DNA metabarcoding of benthic samples compared to eDNA metabarcoding of streamwater. Top row: family taxonomic level. Middle row: genus taxonomic level. Bottom row: species taxonomic level.

**Figure 5.**
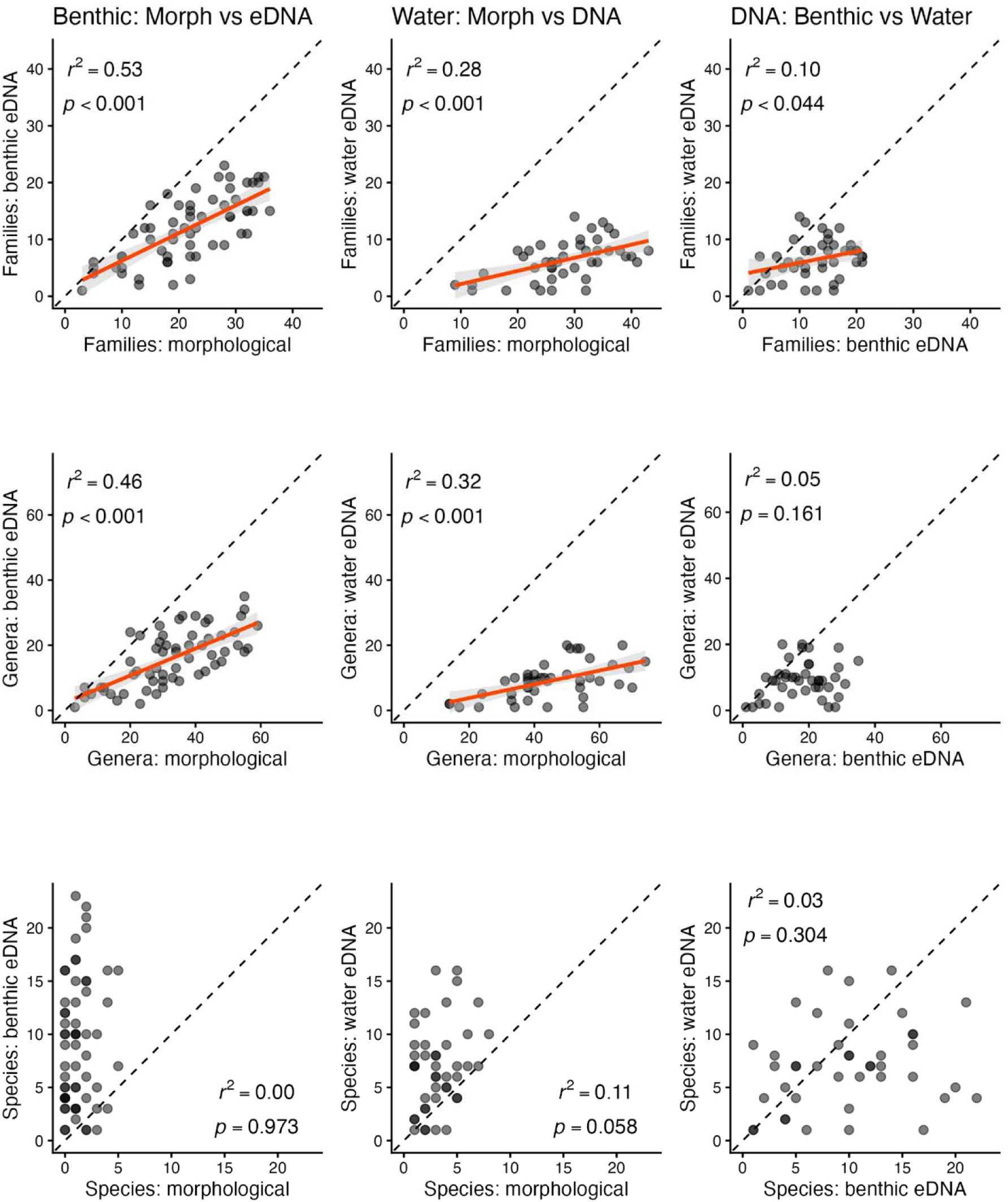
Linear regressions of pairwise methods comparisons of richness measured across taxonomic levels. Dashed lines represent a 1:1 relationship. Left column: DNA morphological of benthic samples (x) compared to metabarcoding comparison of benthic samples (y). Middle column: morphological comparison of benthic + edge samples (x) compared to eDNA metabarcoding of streamwater (y). Right column: DNA metabarcoding of benthic samples (x) compared to eDNA metabarcoding of streamwater (y). Top row: family taxonomic level. Middle row: genus taxonomic level. Bottom row: species taxonomic level. Red solid lines represent statistically significant linear regressions (p < 0.05).

**Table 1.** Results of GLMMs testing for significant deviations of log-response ratios from zero of methods comparisons at different taxonomic levels. Data are estimates of mean log-response ratios from the GLMM, followed by test statistics and p-values in parentheses. The sign of log-response ratio indicates the direction of the difference in richness from the first method relative to the second, and the magnitude of the log-response ratio indicates the magnitude of that difference.

| Methods Comparison | Family | Genus | Species |
| --- | --- | --- | --- |
| In(Metabarcoding-benthic/<br>Morphological-benthic) | -0.641<br>$t_{6.01} = -5.36$<br>$p = \mathbf{0.002}$ | -0.751<br>$t_{5.914} = -5.73$<br>$p = \mathbf{0.001}$ | 1.57<br>$t_{6.382} = 5.03$<br>$p = \mathbf{0.002}$ |
| In(Metabarcoding-water/Morphological-benthic+edge) | -1.69<br>$t_{5.89} = -8.83$<br>$p < \mathbf{0.001}$ | -1.85<br>$t_{5.827} = -8.55$<br>$p < \mathbf{0.001}$ | 0.664<br>$t_{4.643} = 3.27$<br>$p = \mathbf{0.038}$ |
| In(Metabarcoding-water/Metabarcoding-benthic) | -0.783<br>$t_{5.60} = -3.62$<br>$p = \mathbf{0.013}$ | -0.808<br>$t_{5.587} = -2.90$<br>$p = \mathbf{0.021}$ | -0.321<br>$t_{4.96} = -1.39$<br>$p = 0.224$ |

**Table 2.** Results of linear regressions testing for significant relationships between taxonomic richness measurements between methods.

| Methods Comparison | Family | Genus | Species |
| --- | --- | --- | --- |
| y=metabarcoding-benthic,<br>x=morphological-benthic | $F_{1.58} = 62.77$<br>$p < \mathbf{0.001}$<br>$r^2 = 0.52$ | $F_{1.58} = 53.39$<br>$p < \mathbf{0.001}$<br>$r^2 = 0.48$ | $F_{1.58} = 0.983$<br>$p < 0.973$<br>$r^2 = 0.00$ |
| y=metabarcoding-water,<br>x=morphological-benthic+edge | $F_{1.42} = 16.93$<br>$p < \mathbf{0.001}$<br>$r^2 = 0.29$ | $F_{1.42} = 19.70$,<br>$p < \mathbf{0.001}$<br>$r^2 = 0.32$ | $F_{1.32} = 3.29$<br>$p < 0.076$<br>$r^2 = 0.08$ |
| y=metabarcoding-water,<br>x=metabarcoding-benthic | $F_{1.40} = 4.43$<br>$p < \mathbf{0.042}$<br>$r^2 = 0.10$ | $F_{1.40} = 2.33$<br>$p < 0.135$<br>$r^2 = 0.06$ | $F_{1.38} = 1.31$<br>$p < 0.197$<br>$r^2 = 0.04$ |

NMDS ordinations by site at the family and genus level (Figure 6), produced a stable 3-axis solution for family (stress = 0.155) and a stable 2-axis solution for genus (stress = 0.178). Multi-response permutation procedure (MRPP) indicated significant differences in taxonomic composition among methods for family (A = 0.037, p < 0.001) and genus (A = 0.028, p < 0.001). Of the 142 families, 372 genera, and 263 species identified (Tables S2-S4), 38 families, 60 genera, and 1 species were common across the four methods (Figure 7). Metabarcoding methods had the largest number of unique taxa (Table 3, Table S5), with benthic DNA metabarcoding having 8 unique families, 30 unique genera, and 94 unique species and eDNA metabarcoding having 17 unique families, 35 unique genera, and 60 unique species. Morphological-Benthic+Edge also had unique taxa, with 8 unique families, 24 unique genera, and 12 unique species. Morphological-Benthic had no taxa unique to that method alone at any taxonomic level.

**Figure 6.**
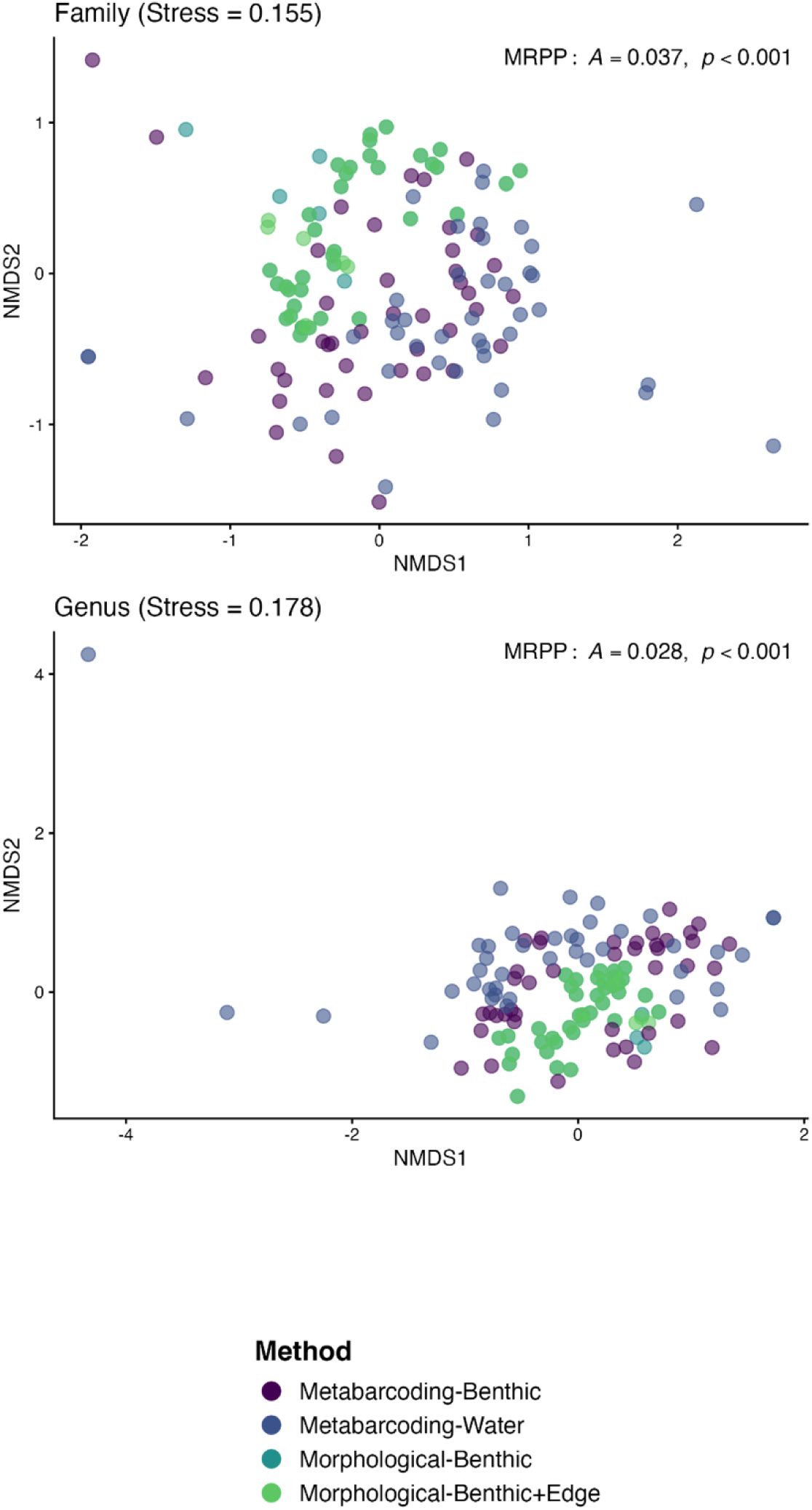
Non-metric multidimensional scaling (NMDS) ordinations of site x taxon matrices at the family and genus taxonomic levels using presence/absence data for the most common taxa (relative abundance > 1%). NMDS ordinations at the species level did not converge on a stable solution. Multiple Response Permutation Procedure (MRPP) results indicate significant differences among methods at both family and genus levels. The family NMDS ordination required a 3-axis solution; here we present the first 2-axes for brevity and present the visualization for the 3-axis solution in the supplementary information.

**Figure 7.**
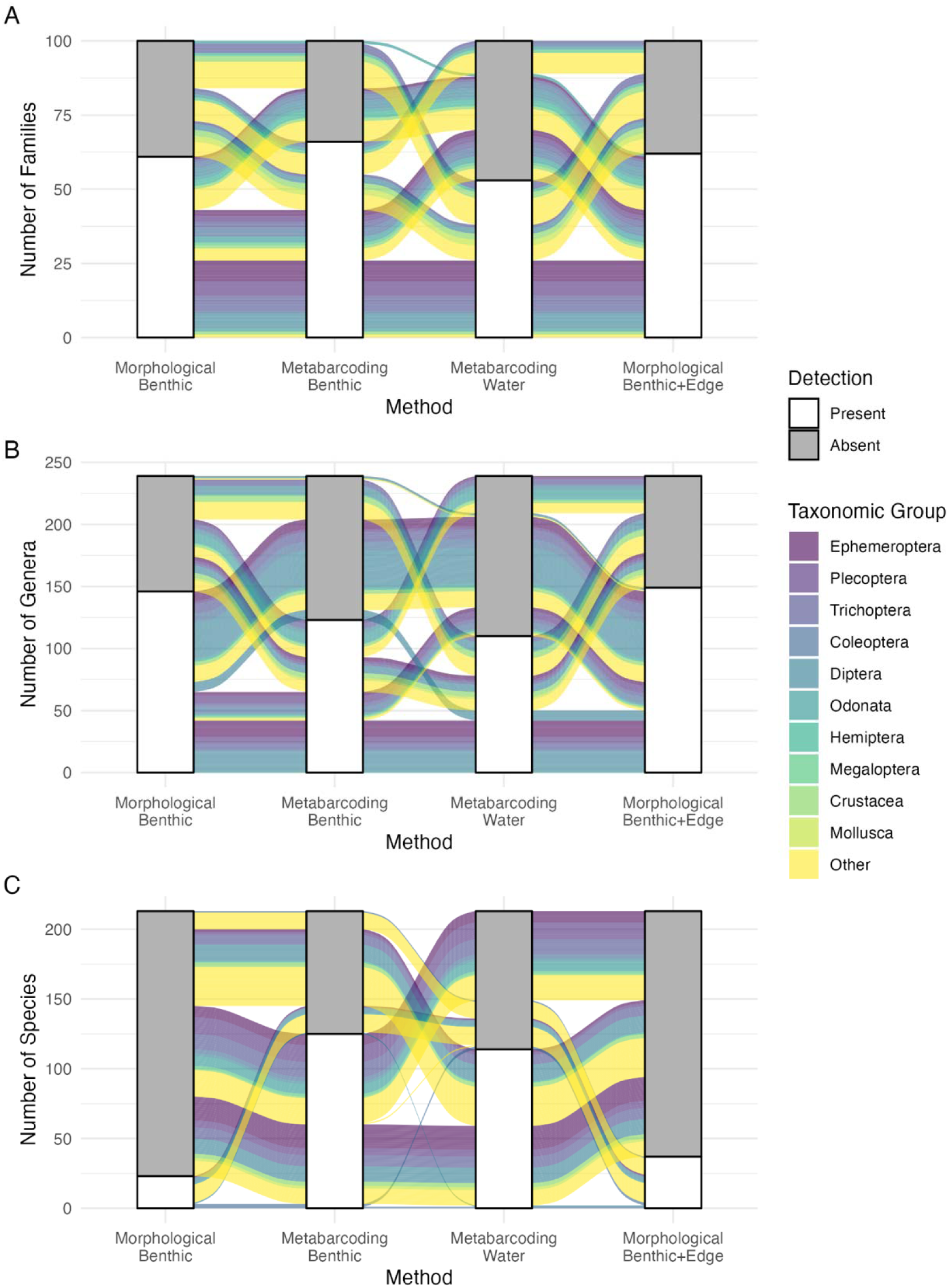
Alluvial plots showing shared taxa among the four different methods for families (A), genera (B), and species (C). Stacked bars indicate taxa present (white) or absent (gray), and flows between the stacked bars connect the same taxa. Due to the large number of families (100) and genera (239), individual taxa are color coded by their broader taxonomic groups. Major insect orders: Ephemeroptera, Plecoptera, Trichoptera, Coleoptera, Diptera, Odonata, Hemiptera, and Megaloptera; Crustacea, Mollusca, and all “Other” families (Annelida, Arachnida, Nematoda, etc.

**Table 3.** Summary of unique taxa found by each method (see Table S4 for the full list of taxa unique to each method).

| Method | Unique Families | Unique Genera | Unique Species |
| --- | --- | --- | --- |
| Morphological-Benthic | 0 | 0 | 0 |
| Morphological-Benthic+Edge | 8 | 24 | 12 |
| Metabarcoding-Benthic | 8 | 30 | 94 |
| Metabarcoding-Water | 17 | 35 | 60 |

## Discussion

Effective freshwater biomonitoring depends on accurate and comprehensive biodiversity assessment, yet the methods available for stream macroinvertebrate identification differ substantially in the taxa they detect, the taxonomic resolution they achieve, and the spatial scales at which they can realistically be deployed. As DNA-based metabarcoding approaches become increasingly integrated into biomonitoring programs worldwide, understanding how they compare to traditional morphological identification, and how different metabarcoding approaches compare to each other, is critical for making informed decisions about method choice and interpreting biodiversity data across studies that use different approaches. Our investigation into this matter revealed three intriguing patterns. First, which method detected more taxa shifted with taxonomic resolution: morphological methods detected more families and genera, while metabarcoding methods detected more species, only partially supporting our first hypothesis. Second, metabarcoding of bulk benthic samples detected more families and genera than water eDNA metabarcoding, but the two methods did not differ significantly for species. Third, and perhaps most striking, the four methods detected largely distinct suites of taxa, with both metabarcoding methods yielding substantially more unique taxa than either morphological method, and with the two metabarcoding methods themselves differing considerably in the taxa they uniquely detected. These results support our second hypothesis. Overall, these results demonstrate that multiple methods used together can describe the most biodiversity, highlighting the need for more work to understand how to use these methods in tandem.

The shift in which method detected more taxa as taxonomic resolution increased contrasts somewhat with the findings of (Emmons et al. 2023), who compared morphological identification to benthic DNA metabarcoding across continental-scale National Ecological Observatory Network sites. (Emmons et al. 2023) found that metabarcoding and morphological methods identified similar levels of alpha diversity on a per sample basis for orders, families, and genera, but that metabarcoding methods detected more unique taxa overall and showed higher gamma diversity at these taxonomic levels. However, (Emmons et al. 2023) did not examine species-level detections, in part because morphological identification of aquatic macroinvertebrate larvae is often limited to genus for many taxa, and species-level identification is frequently impossible for larval stages of many groups (Haase et al. 2010, Elbrecht et al. 2017). Our more conservative thresholds for retaining high-quality reads (99% confidence for species-level assignments) likely reduced the number of spurious species-level identifications relative to other studies, but also likely reduced the number of taxa we identified using metabarcoding methods relative to studies using less conservative thresholds.

That morphological methods identified more families and genera, while metabarcoding detected more species, likely reflects fundamental differences in how each method samples biodiversity. In morphological identification, all organisms in the sample are visually sorted and identified, meaning common, abundant, high-biomass taxa are reliably detected because they are physically present in large numbers. In contrast, metabarcoding involves sequential subsampling steps: DNA extraction, PCR amplification, and sequencing; each of which represents a subsample of the previous step (Elbrecht and Leese 2015, Martins et al. 2021). Common, high-biomass taxa may therefore be diluted relative to their true abundance in the sample, reducing the probability that their reads dominate the sequencing output. Further, an additional consideration is that PCR primers show taxonomic biases in amplification and work better for some taxa than others (Clarke et al. 2014, Elbrecht and Leese 2015), so this bias could influence the representation of high-biomass taxa if primers do not effectively amplify their DNA. Conversely, metabarcoding can detect rare or cryptic taxa present at very low abundances from trace DNA, including species within genera whose larvae cannot be distinguished morphologically, taxa that would be easily overlooked or unidentifiable during visual sorting.

The greater detection of families and genera by bulk benthic metabarcoding compared to water eDNA metabarcoding is consistent with the expectation that the DNA concentration of target taxa is generally higher in extracted tissue from the benthic sample than in filtered water, where organismal DNA is diluted and degraded (Deiner et al. 2016, Hajibabaei et al. 2019a). This “watered-down biodiversity” effect has been documented in other systems comparing bulk benthic and water eDNA (Macher et al. 2018, Hajibabaei et al. 2019a, van der Lee et al. 2026), though those studies were limited to single watersheds or small numbers of sites and did not include morphological comparisons. Our results confirm and extend this pattern to a large-scale, multi-watershed context, providing stronger evidence that it is a generalizable phenomenon rather than a site-specific artifact. The absence of a significant difference in total species richness between the two metabarcoding methods likely reflects that both methods excel in the species level identifications that morphological identifications do not, though we found differences in the identity of the species that the two metabarcoding methods detected.

Perhaps the most ecologically interesting finding of our study was the substantial difference in which taxa were uniquely detected by benthic versus water eDNA metabarcoding. Taxa uniquely detected by water eDNA were dominated by soft-bodied organisms: oligochaetes (Nais, Dero, Limnodrilus), leeches (Helobdella), and earthworms (Lumbricus terrestris, Amynthas). Additionally, water eDNA detected taxa that are more associated with the water column or water surface, like copepods and water striders (Aquarius), but also unionid mussels that filter large amounts of water. In contrast, taxa uniquely detected by benthic eDNA were overwhelmingly hard-bodied EPT insects, including 18 Ephemeroptera species, 17 Plecoptera species, and 14 Trichoptera species, as well as Odonata and Coleoptera. This pattern strongly suggests that water eDNA dynamics are fundamentally shaped by organismal physiology. Soft-bodied organisms may shed DNA more readily into the water column, while hard-bodied arthropods with exoskeletons may shed relatively little DNA into water, perhaps instead contributing DNA primarily through molting events that leave genetic material concentrated in the substrate (Andruszkiewicz Allan et al. 2021, Trimbos et al. 2021, Lancaster et al. 2025). Additionally, riparian taxa detected in water samples (earthworms, spiders, etc) likely entered the water via surface runoff and precipitation, a phenomenon documented in other stream eDNA studies (Deiner et al. 2016, Reves et al. 2026). The presence of these taxa in water eDNA samples is an important consideration for bioassessment programs, particularly cutting-edge approaches that rely on automated water sampling with remotely controlled instruments. Water sampling introduces non-target signals that could complicate interpretation of community composition data, and also may not adequately sample target organisms effectively if target organisms do not shed much eDNA.

Our results collectively highlight that the four methods each have distinct biases that favor detection of particular components of the macroinvertebrate community. Morphological benthic sampling excels at detecting abundant, readily identifiable taxa — particularly EPT insects that dominate benthic kicknet samples — but is constrained to genus level for most groups and provides less information on soft-bodied, rare, or morphologically cryptic taxa (Haase et al. 2010, Elbrecht et al. 2017). Adding stream-edge sampling broadens taxonomic coverage slightly, capturing taxa in riparian-adjacent microhabitats that benthic sampling misses (Gill et al. 2024), though the unique taxa detected by this method (8 families, 24 genera, 12 species) were modest relative to unique taxa identified from the metabarcoding methods. Bulk benthic DNA metabarcoding captures a similar taxonomic footprint to morphological methods at family and genus levels (Emmons et al. 2023, Múrria et al. 2024), with the added advantage of species-level resolution for hard-bodied EPT insects, but its processing is still tied to the physical sample collected and may miss soft-bodied taxa that are rare in or absent from the benthic zone. Water eDNA metabarcoding, while capturing fewer families and genera overall, uniquely detects soft-bodied and planktonic taxa (Hajibabaei et al. 2019a), providing a distinct biodiversity signal that none of the other methods recover, though it is more susceptible to non-target signals from terrestrial and riparian organisms. If resources allow only one method, the choice should be guided by the goals of the bioassessment and the taxonomic resolution required: morphological methods remain the gold standard for genus-level and family composition assessments of hard-bodied benthic invertebrates, while metabarcoding of benthic samples provides broader taxonomic coverage including species-level identification, and water eDNA offers a complementary window into soft-bodied and mobile taxa.

Together, these results underscore that no single method captures a complete picture of stream macroinvertebrate biodiversity, and that choices about sampling and identification approach carry meaningful consequences for biodiversity inference. The discordance between eDNA methods in the taxa they detect points to an important and underexplored frontier in stream bioassessment: understanding the eDNA production, transport, and decay dynamics of specific taxa. The biology of eDNA shedding, determined by organism physiology, behavior, abundance, and life history; fundamentally shapes what is and is not detectable in a water sample, yet these processes remain poorly characterized for most freshwater invertebrates (Andruszkiewicz Allan et al. 2021, Trimbos et al. 2021). Once shed, eDNA undergoes ‘spiraling’ in stream systems, cycling between transport downstream by stream flow, deposition into sediments, and resuspension back into the water column; while simultaneously subject to degradation by UV radiation and microbial activity (Shogren et al. 2017, 2019, Harrison et al. 2019). These processes collectively determine whether a taxon’s DNA is detectable at a given sampling location and time, and vary considerably across taxa, stream conditions, and seasons. Our results suggest that hard-bodied EPT insects, which are among the most ecologically important and widely used bioindicators in freshwater bioassessment, are systematically underrepresented in water eDNA relative to their actual occurrence. Until these dynamics are better understood across a broader range of taxa and stream types, combining multiple sampling approaches will remain the most reliable strategy for comprehensive macroinvertebrate biodiversity assessment in streams.

## Author contributions (CRediT taxonomy)

Conceptualization: DCA, ZGC, LGL, KTH, CS, FI, LEVS, KC, AS, CLA, MTB, TN, SE, MCM, YH

Data curation: DCA, ZGC, LGL, CS, FI, LEVS, KC, AS, CLA, SE, TA, BAG

Formal analysis: DCA, ZGC, LGL, CS, FI, LEVS, KC, AS, CLA, SE

Funding acquisition: DCA, AR, MTB, TN, MCM, YH

Investigation: DCA, ZGC, KTH, CS, LEVS, KC, AR, AS, CLA, KL, MTB, MM, SCC, SS, MCM, TA, MHB, BAG, VS, RM

Methodology: DCA, ZGC, LGL, CS, FI, LEVS, KC, AR, AS, CLA, MTB, SE, MM, SCC, MCM, TA, BAG, VS

Project administration: DCA, ZGC, AR, AS, CLA, KL, MTB, SS, MCM, TA, BAG, VS

Software: DCA, ZGC, CS, SE

Resources: DCA, ZGC, CS, LEVS, KC, AS, CLA, KL, MTB, SCC, SS, MCM, BAG

Supervision: DCA, ZGC, AR, AS, CLA, KL, MTB, TN, SS, MCM, TA, BAG

Validation: DCA, ZGC, CS, TA, BAG

Visualization: DCA, ZGC, KTH, CS, LEVS, KC, SE

Writing – original draft: DCA, ZGC, LGL, KTH, CS, FI, LEVS, KC

Writing – review & editing: all co-authors

## Data Availability Statement

All data and R code used in this study are publicly available at https://doi.org/10.5281/zenodo.22943991 and https://github.com/dcallen/StreamCLIMES. These data follow the FAIR (Findable, Accessible, Interoperable, and Reusable) data principles.

## Funding statement

Funding for this project was provided by the US National Science Foundation, as part of the collaborative StreamCLIMES (awards: 1802714, 1802766, 1802811, 2150626, 2207680) and DISTANCe (awards: 2406856, 2406857, 2406858) projects. This work was also supported by the USDA National Institute of Food and Agriculture and Hatch Appropriations under Project #PEN04817 and Accession #7003724.

## Conflicts of interest disclosure

The authors declare no conflicts of interest.

## Supplementary Figures and Tables

**Figure S1.**
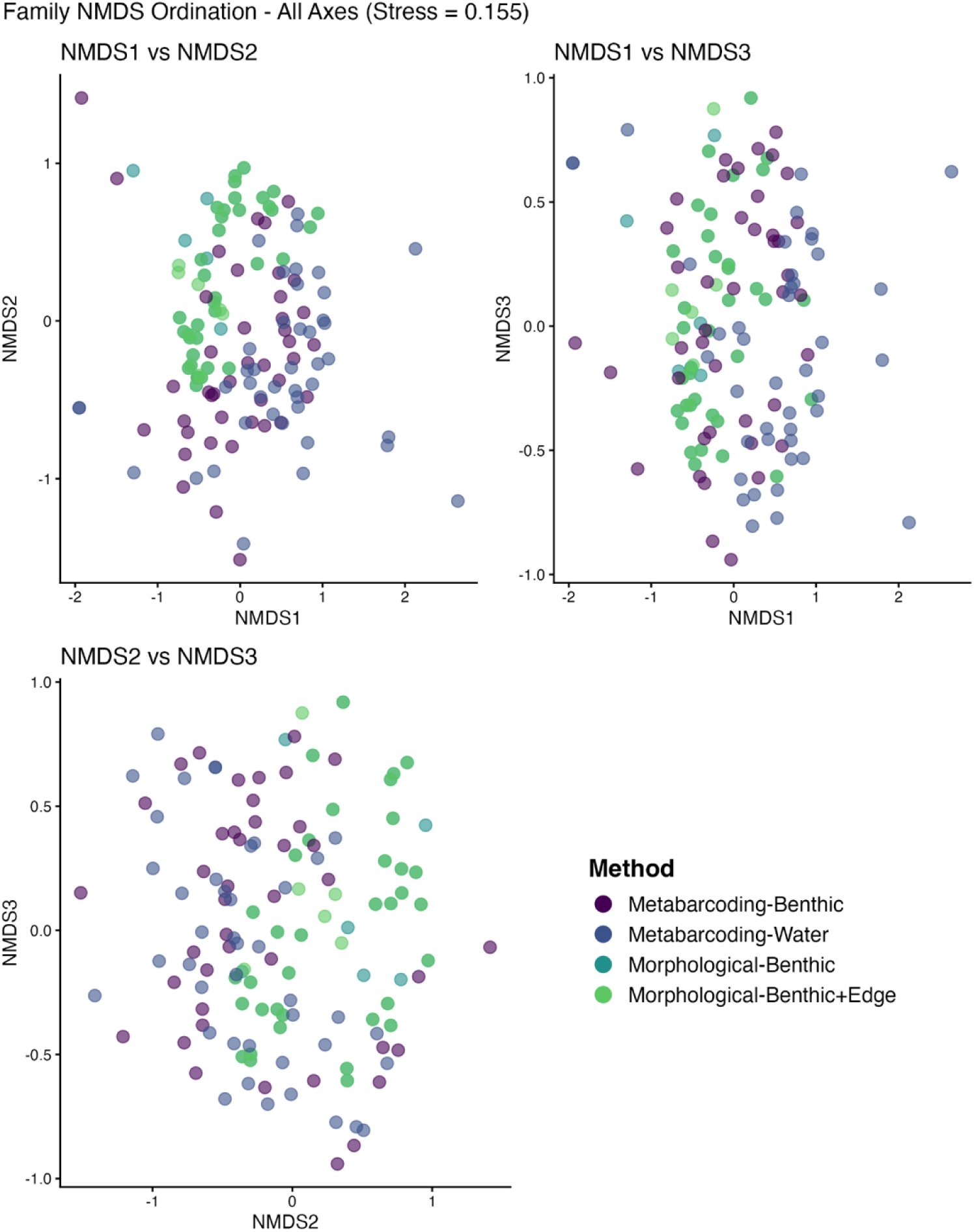
Non-metric multidimensional scaling (NMDS) ordination of site x taxon matrices at the family taxonomic level using presence/absence data for the most common taxa (relative abundance > 1%).

**Table S1.** Sampling site locations and method coverage for all sites where at least one morphological and one metabarcoding method were used (n = 62). Basin codes: BLUE = Blue River (Oklahoma), ICHA = Ichawaynochaway Creek (Georgia), KING = Kings Creek (Kansas), MAYF = Mayfield Creek (Alabama), PASS = Passage Creek (Virginia), SNMT = Sierra Nevada Mountain Tributaries (California), WPLP = West Prong Little Pigeon River (Tennessee). Flow Class indicates whether the site was perennial (continuously flowing) or intermittent (periodically dry) during the study period. X indicates that data from that method were collected and used in analyses at that site. Morphological-Benthic: traditional morphological identification of benthic kicknet samples; Morphological-Benthic+Edge: morphological identification of combined benthic and stream-edge samples; Metabarcoding-Benthic: DNA metabarcoding of homogenized bulk benthic samples; Metabarcoding-Water: eDNA metabarcoding of filtered streamwater samples.

TableS1_Paired_Sites.csv

**Table S2.** Detection of macroinvertebrate families across four sampling and identification methods. Presence (1) and absence (0) of each family is indicated for Morphological-Benthic, Morphological-Benthic+Edge, Metabarcoding-Benthic, and Metabarcoding-Water methods. Families are organized by taxonomic order and sorted alphabetically within each order. TableS2_Family_Taxa_Table.csv

**Table S3.** Detection of macroinvertebrate genera across four sampling and identification methods. Presence (1) and absence (0) of each genus is indicated for Morphological-Benthic, Morphological-Benthic+Edge, Metabarcoding-Benthic, and Metabarcoding-Water methods. Genera are organized by taxonomic order and sorted alphabetically within each order. TableS3_Genus_Taxa_Table.csv

**Table S4.** Detection of macroinvertebrate species across four sampling and identification methods. Presence (1) and absence (0) of each species is indicated for Morphological-Benthic, Morphological-Benthic+Edge, Metabarcoding-Benthic, and Metabarcoding-Water methods. Species are organized by taxonomic order and sorted alphabetically within each order. TableS4_Species_Taxa_Table.csv

**Table S5.** Taxa detected exclusively by a single sampling and identification method across all study sites. Rows represent individual taxa organized by taxonomic level (Family, Genus, Species) and sorted alphabetically within each taxonomic group. Columns indicate which method uniquely detected each taxon, where “X” indicates detection and blank cells indicate non-detection. Morphological-Benthic is omitted as no taxa were detected exclusively by this method. Taxa are further organized by broader taxonomic group (major insect orders: Ephemeroptera, Plecoptera, Trichoptera, Coleoptera, Diptera, Odonata, Hemiptera, and Megaloptera; Crustacea; Mollusca; and Other).

TableS5_Unique_Taxa_Table.csv

## Notes

https://github.com/dcallen/StreamCLIMES

https://doi.org/10.5281/zenodo.22943991

### Competing Interest Statement

The authors have declared no competing interest.

